# Local CpG density affects the trajectory of age-associated epigenetic changes

**DOI:** 10.1101/2021.07.08.451539

**Authors:** Jonathan Higham, Qian Zhang, Rosie M. Walker, Sarah E. Harris, David M. Howard, Emma L. Hawkins, Anca-Larisa Sandu, J. Douglas Steele, Gordon D. Waiter, Alison D. Murray, Kathryn L. Evans, Andrew M. McIntosh, Peter M. Visscher, Ian J. Deary, Simon R. Cox, Duncan Sproul

## Abstract

DNA methylation is an epigenetic mark associated with gene repression and genome stability. Its pattern in the genome is disrupted with age and these changes can be used to statistically predict age with epigenetic clocks. Rates of aging inferred from these clocks correlate with human health. However, the molecular mechanisms underpinning age-associated DNA methylation changes are unknown. Local DNA sequence plays a strong role in programming DNA methylation levels at individual loci independently of age, but its influence on age-associated DNA methylation changes is unknown. We analysed longitudinal human DNA methylation trajectories at 345,895 CpGs from 600 individuals aged between 67 and 80 to understand the factors responsible for age-associated epigenetic changes at individual CpGs in the genome. We show that changes in methylation with age are especially apparent at 8,322 low CpG density loci. Using SNP data from the same individuals we demonstrate that DNA methylation trajectories are affected by local sequence polymorphisms at 1,487 loci with low CpG density. More generally, we find that local CpG density is a strong determinant of a CpG’s methylation trajectory and that CpGs located in low CpG density regions are particularly prone to change. Overall, our results demonstrate that local DNA sequence influences age-associated DNA methylation changes in humans *in vivo*. We suggest that this occurs because interactions between CpGs reinforce maintenance of methylation patterns in CpG dense regions.

## Introduction

DNA methylation is the most common DNA modification found in mammals and is a repressive epigenetic mark observed predominantly at cytosines in CpG dinucleotides. In mammals, it is largely erased from the somatic genome in early development and re-established later by the *de novo* DNA methyltransferases 3A and 3B (DNMT3A and DNMT3B)^1^. This wave of *de novo* methylation results in a pervasive methylation landscape where 70-80% of CpGs are methylated in most human tissues^2^. Short regions lacking DNA methylation often correspond to promoters and other regulatory elements particularly CpG islands (CGIs)^3^. Subsequently DNA methylation is largely maintained by the action of the maintenance DNA methyltransferase, DNMT1^4^.

Despite maintenance methylation, the DNA methylation landscape alters with age. Overall, the DNA methylation content of the genome reduces with age^5^ but individual loci gain methylation. Changes at some loci are sufficiently reproducible across individuals as to enable the statistical derivation of accurate predictors of age termed epigenetic clocks^6,7^. Furthermore, deviations between epigenetic clock predicted age and true chronological age have been associated with human health^8^. Similar epigenetic clocks have been derived for other mammalian species^9–12^. In mice, interventions associated with increased lifespan also associate with decreases in epigenetic age measured by murine epigenetic clocks^9,11^. In addition to epigenetic clocks, variable DNA methylation changes between individuals termed epigenetic drift has also be described^13^. For example, increased divergence in the DNA methylation patterns of twins is observed with age^14^.

Losses of DNA methylation with age are thought to occur primarily in heterochromatic, late replicating genomic regions^15^. Conversely, DNA methylation gains have been associated with CGIs that are targeted by polycomb repressive complexes in embryonic stem (ES) cells^16–18^. However, the molecular mechanisms underpinning all age-associated DNA methylation alterations remain unknown. Analysis of methylation in human populations independent of age has shown that differences in the methylation level of individual loci between individuals frequently associate with sequence polymorphisms. These have been characterised as allele-specific methylation events and methylation quantitative trait loci (meth-QTLs)^19,20^. This suggests that DNA sequence can program local DNA methylation levels, a hypothesis supported by the inheritance pattern of allele-specific methylation in families^21^ and analysis of the methylation patterns of ectopic DNA sequences integrated into cell lines^22^. Whether DNA sequence plays a role in age associated changes in DNA methylation is less clear. Some studies have provided evidence that genetic variants affect how DNA methylation changes with age at individual loci^23,24^. However, a study of mice possessing a copy of human chromosome 21 suggested that local sequence plays little role in determining the rate of age-associated DNA methylation changes^25^. In this study the human chromosome accumulated age-associated changes a similar rate to the mouse genome rather than the rate observed in its native human context.

Here we examine longitudinal DNA methylation patterns at 345,895 individual CpGs in blood DNA samples from people aged between 67 and 80 to understand the mechanisms that underpin age-associated epigenetic changes. We demonstrate a strong relationship between DNA methylation changes with age and local CpG density.

## Results

### Longitudinal methylation trajectories reveal changes at individual epigenetic clock loci

In order to understand the factors that are responsible for age-associated alterations in DNA methylation at the CpG level, we analysed longitudinal DNA methylation data collected from blood samples taken from the Lothian Birth Cohort of 1936 (LBC)^26–28^. This cohort consists of 1,091 individuals, whose blood was assayed at multiple time-points on Illumina Infinium 450k arrays between the ages of 67 and 80 (see *Table 1*). To robustly quantify DNA methylation alterations with age, we focused on the 600 individuals for whom 3 or more datapoints were available and 345,895 reliably measured autosomal CpG probes whose signal is not directly affected by SNPs or cross-hybridisation^29^.

**Table 1.** Demographics of participants used in this study. The mean age in years at each measurement are indicated along with the range. The number of observations at each measurement are also indicated. In total 351 individuals had data for all 4 measurements and 249 for three of the measurements.

|  | 1<sup>st</sup> measurement | 2<sup>nd</sup> measurement | 3<sup>rd</sup> measurement | 4<sup>th</sup> measurement |
| --- | --- | --- | --- | --- |
| mean age<br>(min/max) | 69.6<br>(67.7, 71.3) | 72.6<br>(71.0, 74.2) | 76.3<br>(74.7, 77.7) | 79.3<br>(78.0, 80.9) |
| no. females | 265 | 271 | 259 | 224 |
| no. males | 295 | 307 | 292 | 238 |

We modelled methylation trajectories for each CpG and individual as linear models of the Infinium beta values with age (example shown in *Figure 1a*). Estimated mean individual rates of change in methylation with age (hereafter slopes), correlated highly with those calculated from the whole cohort in a cross-sectional manner suggesting they are a robust measure (*Supplementary Figure 1a*, Pearson’s R = 0.988, p < 2.2×10^−16^). We first examined the CG probe *cg16867757* in the *ELOV2L* promoter which shows strong age-associated changes in DNA methylation to test whether our approach could measure age-associated changes at individual loci^7,30^. We observed a highly significant gain of methylation with age at the locus in the LBC cohort, validating our approach (*Figure 1a*, T-test, p < 2.2×10^−16^).

**Figure 1.**
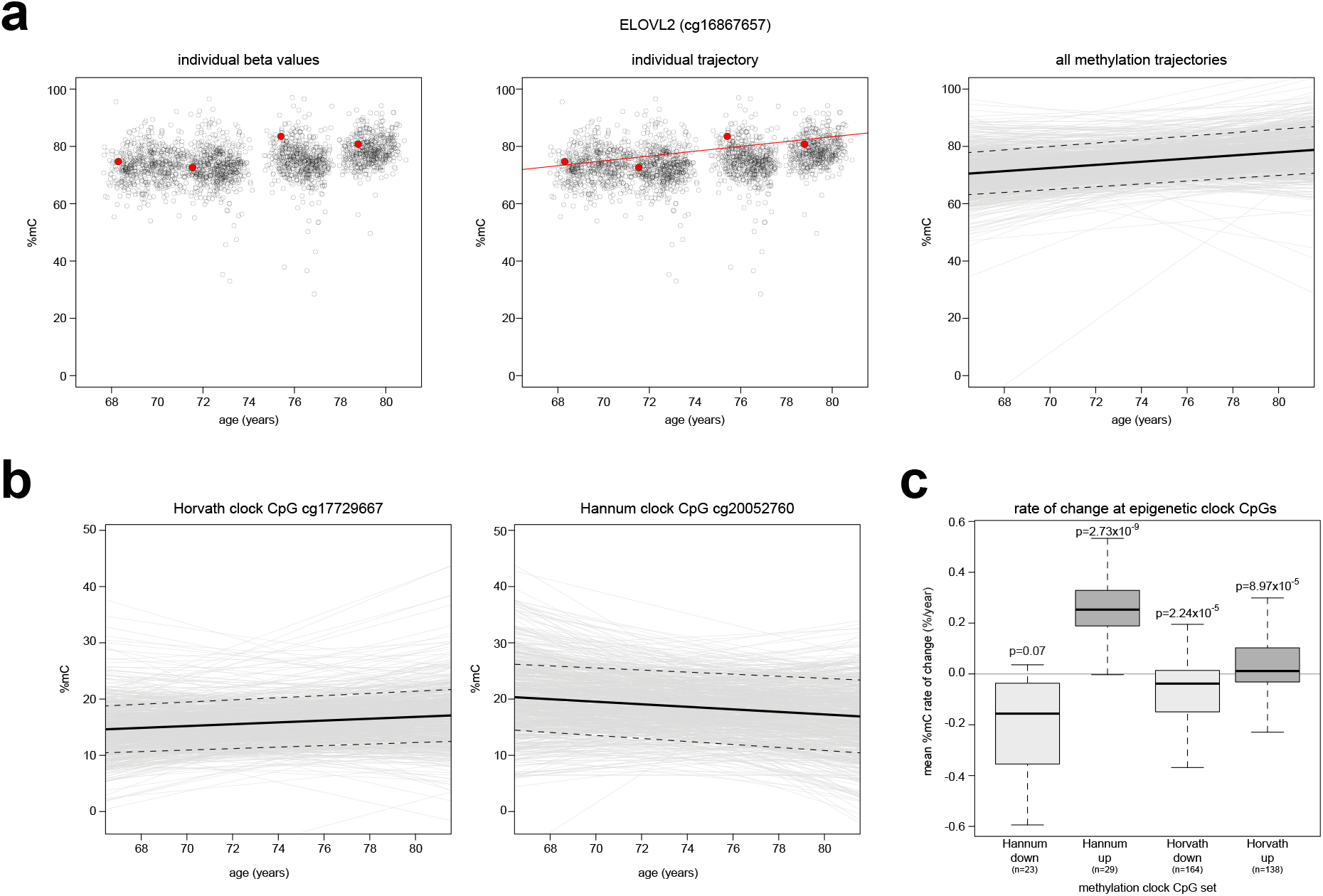
Epigenetic clock CpGs show consistent changes in methylation with age in individuals. **a)** ELOLV2 shows increases in methylation across individuals. Plots of methylation levels at ELOVL2 CpG cg16867657 showing data from one individual in the cohort and their methylation trajectory (left and middle panel, red points and line), and methylation trajectories for all individuals (right panel, grey lines). In the right panel, the bold line indicates the mean methylation trajectory, and the dashed lines are the 95% confidence intervals. **b)** Examples of methylation trajectories observed for epigenetic clock CpGs. Individual methylation trajectories are indicated by grey lines. The mean methylation trajectory is indicated by the bold line and the dashed lines are the 95% confidence intervals. **c)** Methylation trajectories recapitulate the predicted behaviour of epigenetic clock CpGs. Boxplots showing the calculated mean rates of change for CpGs that are part of the Hannum or Horvath epigenetic clocks split by their reported direction of change. P-values were calculated using T-test. Lines=median; Box=25th–75th percentile; whiskers=1.5× interquartile range from box; n indicates the number of CpGs in each group.

To further test whether changes at individual loci could be reliably measured, we examined epigenetic clock loci. Although epigenetic alterations with age have been widely quantified using epigenetic clocks, the longitudinal behaviour of individual epigenetic clock loci remains unexplored. Given their use in predicting age, they would be expected to show consistent trajectories between individuals. We therefore assessed the behaviour of the CpGs that are included in the widely used Hannum and Horvath epigenetic clocks^6,7^. 88% CpGs in the Hannum epigenetic clock and 80% of the CpGs in the Horvath epigenetic clock had a statistically significant change with age in the direction predicted by the original studies (46 out of 52 and 241 out of 302 respectively, p < 0.05, T-Test, examples in *Figure 1b* and aggregate analysis *Figure 1c*)^6,7^. The slopes of the CpGs making up these epigenetic clocks were also consistent with those calculated from a cross-sectional cohort of 5,101 individuals from the Generation Scotland study who had their DNA methylation levels profiled on Illumina EPIC arrays (*Supplementary Figure 1b*)^31,32^. The absolute rate of change of clock CpGs were modest compared to other CpGs whose methylation levels changed significantly with age in accordance with observations of other epigenetic clocks (*Supplementary Figure 1c*)^11^.

Our analyses suggest that individual methylation trajectories in the LBC cohort can measure changes at individual CpGs known to display age-associated alterations in DNA methylation levels.

### A subset of CpGs gain methylation in later life

Having tested that we can measure predicted changes in DNA methylation with age, we then examined the rates of change at individual CpGs across the genome.

Of the 345,895 CpGs in the dataset, 182,760 CpGs (52%) had a significant change in DNA methylation with age (Bonferroni corrected p < 0.01, T-test of individual linear model slopes). The distribution of mean slopes for these CpGs was significantly skewed towards loci gaining DNA methylation (*Figure 2a*, p<2.2×10^−16^ by T-test of mean linear model slopes). A distinct shoulder of CpGs with more rapid gains of methylation was also apparent on the histogram. We defined these rapidly gaining CpGs as those with a rate of methylation change > 1.6% per year (8,322 rapid gain CpGs, *Supplementary Table 1*; example *cg22926528* shown in *Figure 2b*). This set of CpGs also showed higher levels of DNA methylation with age in 406 individuals from the Generation Scotland cohort aged >= 65 years (*Figure 2c,* p < 2.2×10^−16^). Our analysis of Generation Scotland also showed that in younger individuals, the same CpGs initially showed lower levels of methylation with age (*Supplementary Figure 2a*, p < 2.2×10^−16^). Rapid gain CpGs also showed significantly higher rates of methylation gain than other CpGs when slopes were corrected for measured white blood cell counts from the LBC cohort (*Supplementary Figure 2b*) suggesting that the observed gains of DNA methylation did not result from altered blood composition with age.

**Figure 2.**
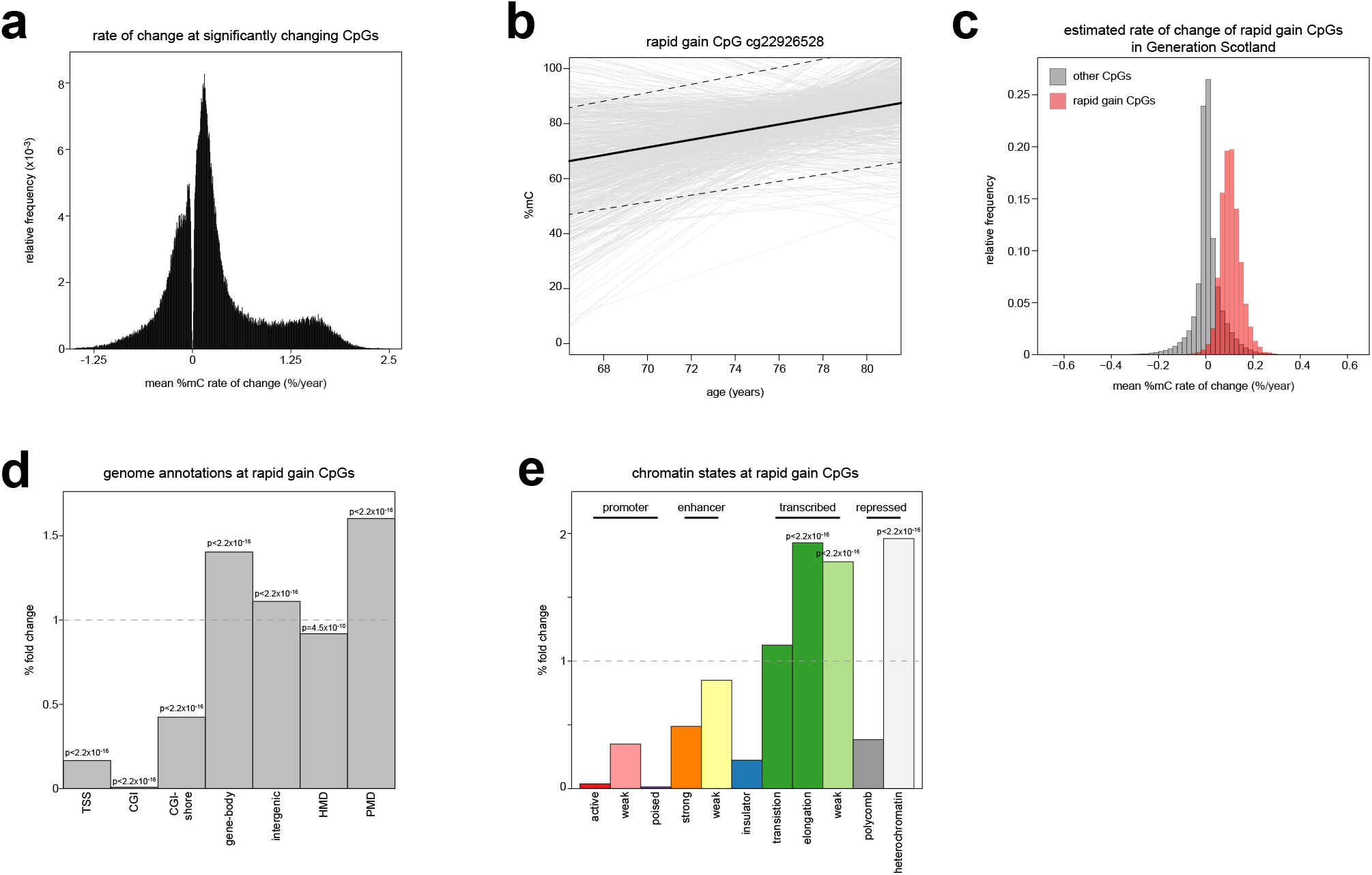
A subset of CpGs gains methylation in later life. **a)** A subset of CpGs gain methylation in LBC. Histogram of the mean methylation trajectories for the 182,760 CpGs whose slope significantly deviates from 0 (T-test, Bonferroni corrected p < 0.01). **b)** Example of a rapid gain CpG. Individual methylation trajectories are indicated by grey lines. The mean methylation trajectory is indicated by the bold line and the dashed lines are the 95% confidence intervals. **c)** Rapid gain CpGs are reproduced in an independent cohort. Histogram of the rate of change in DNA methylation calculated from individuals aged ≥65 from Generation Scotland cohort for rapid gain CpGs (red) and all other CpGs (grey). **d)** Rapid gain CpGs are depleted from CGIs and enriched in gene-bodies and intergenic regions. Barplot showing the % fold change observed for rapid gain CpGs in different genome annotations versus the background of all analysed CpGs. P-values are from 2-sided Fisher’s exact tests. PMDs = partially methylated domains; HMDs = highly methylated domains. **e)** Rapid gain CpGs are enriched in transcription and heterochromatin states in GM12878 cells. Barplot showing the % fold change observed for rapid gain CpGs in different chromatin states in GM12878 cells versus the background of all analysed CpGs. Shown are significant P-values from 1-sided Fisher’s exact tests.

To understand why these CpGs might gain methylation, we examined where they were located in the genome. Compared to all other CpGs in the dataset, the rapid gain CpGs were significantly depleted from CGIs and the regions surrounding CGIs which have been termed shores^33^ (*Figure 2d*). They were instead enriched in the bodies of coding genes (63.3% of CpGs, *Figure 2d*) and large hypomethylated genomic regions termed partially methylated domains (PMDs)^34^ defined across 40 tumour and 9 normal samples (34.6% of CpGs, *Figure 2d*)^15^. PMDs are known to be heterochromatic, gene poor and to have a lower CpG density than other regions of the genome^15,34,35^. Consistent with their enrichment in PMDs, the regions surrounding rapid gain CpGs had a significantly lower CpG density than other CpGs analysed (*Supplementary Figure 2c, Wilcoxon test p* < 2.2×10^−16^).

To further understand the characteristics of this set of CpGs, we cross-referenced them to chromatin state data generated by the ENCODE and Roadmap Epigenomic projects^36,37^. These projects have used hidden Markov models to partition the genome into distinct chromatin states (ChromHMM)^38^. Consistent with their observed enrichment in gene bodies and PMDs, the rapid gain CpGs were significantly enriched in ENCODE-defined transcriptional and heterochromatic chromatin states in GM12878 lymphoblastoid cells (11.0% and 57.7% of CpGs with transcriptional and heterochromatin annotations respectively, *Figure 2e*)^36^. Similarly, they were most enriched in the heterochromatin-associated quiescent state across a set of 23 primary blood cell types whose chromatin states were defined by the Roadmap Epigenomics project (*Supplementary Figure 2d*)^37^. Transcriptional states were also enriched in these primary blood cells but to a lesser degree (*Supplementary Figure 2d*).

Taken together, these observations suggest that changes in methylation in the LBC cohort are most apparent at a subset of CpGs, largely located in heterochromatic, low CpG density regions.

### Local SNPs associate with altered CpG methylation trajectories

Having uncovered a set of CpGs which rapidly gained methylation with age, we then asked what factors led to differences in the trajectories of DNA methylation alterations between CpGs. Previous reports suggest a potential influence of genetic variation on age-associated DNA methylation changes^23,24^ so we tested for associations between local (*cis*; within 1Mb) SNPs and the rate of change of DNA methylation at individual CpGs with age across the cohort.

Our analysis uncovered 4,673 slope-Quantitative Trait Loci (slope-QTLs) representing 1,456 unique CpGs (examples in *Figure 3a*). The linkage disequilibrium structure of the human genome means that linked SNPs would be expected to be associated with the same CpG. In order to determine the number of independent associations, we used conditional analysis to resolve these slope-QTLs into 1,487 SNP-CpG pairs designating the closest independent SNP in each case as the lead SNP (*Supplementary Table 2*). Only 31 CpGs were independently associated with more than 1 SNP. Each lead SNP was significantly associated with the rate of change of methylation at a mean of 1.06 CpGs (range 1 to 16, *Supplementary Table 2*). We validated these associations using another method by testing for an age x genotype interaction effect in a standard linear model. Nearly all of these SNP-CpG pairs (1,334, 90.3%) had a significant interaction (Bonferroni corrected, p<0.05) and the effect size determined from the slope of the individual linear models, and the effect size of the age x genotype interaction within the population were significantly correlated (Spearman Rho = 0.977, p < 2.2×10^−16^, *Supplementary Figure 3a*).

**Figure 3.**
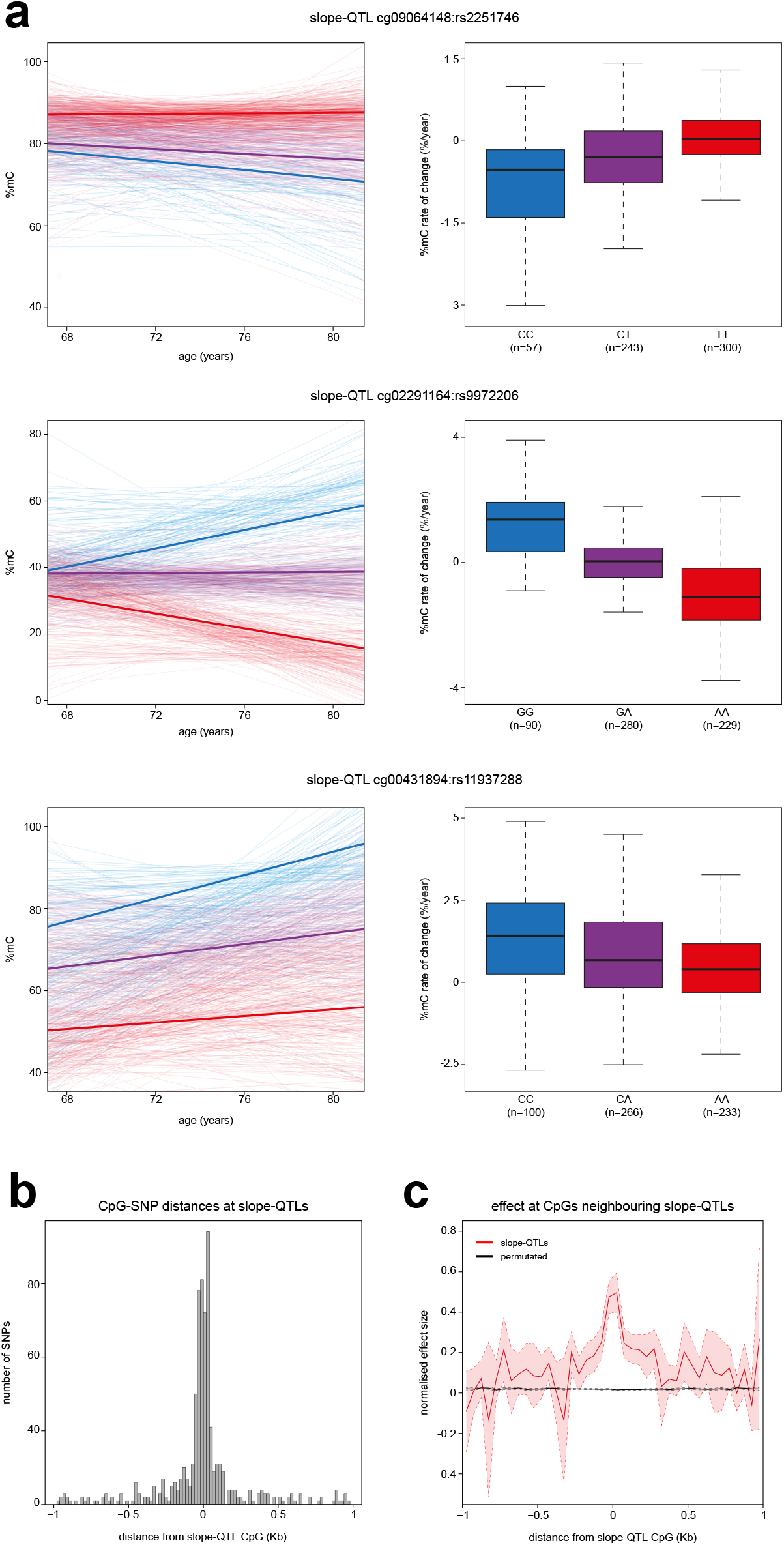
**a)** Examples of slope-QTLs. Spaghetti plots and boxplots of 3 slope-QTL CpG-SNP pairs. Left, spaghetti plots of individual methylation trajectories separated by genotype. Thin lines represent individual methylation trajectories and thick lines the mean methylation trajectory for that genotype. Right, boxplots of slope separated by genotype. Lines=median; Box=25th–75th percentile; whiskers=1.5× interquartile range from box. SNP genotypes are annotated relative to the forward strand. **b)** Slope-QTL SNPs are located in close proximity to the CpGs they affect. Histogram of the distances between slope-QTL lead SNPs and the CpGs they are paired with. **c)** Nearby CpGs are also affected by slope-QTL SNPs. Line plot of the effect sizes calculated for CpGs within −/+ 1Kb of slope-QTL CpGs using each slope-QTL’s lead SNPs. Plotted is the mean normalised effect size in 50bp Windows. Bold lines show the mean effect size and dashed lines and shaded area show the 95% confidence intervals. The data are shown in red and the results of 1000 random permutations shown in black.

We also tested for *trans*-slope-QTLs independent of SNP-CpG genomic distance. This analysis uncovered 9 significant SNP-CpG pairs (*Supplementary Table 3*). The low number was likely due to the multiple testing burden associated with testing every SNP against every CpG. Given the low number of observed *trans*-slope-QTLs, we focused on the analysis of *cis*-slope-QTLs.

Although we had set a threshold of 1Mb when uncovering *cis* slope-QTL, the lead SNPs were located close to the slope-QTL CpGs (*Figure 3b*). At 53% of the slope-QTLs the lead SNP and CpG were within 1Kb of each other. Whereas most SNPs (1346/1397, 96.3%) were only associated with a single CpG at the genome-wide Benjamini-Hochberg corrected significance FDR < 0.05, we observed that other CpGs close to those in slope-QTLs showed correlated effects that were below the multiple-testing corrected significance threshold (*Figure 3c*). This strongly suggests that these slope-QTLs are driven by specific effects of genotype on methylation change with age within local genomic regions. It also makes it unlikely that slope-QTLs are caused by the disruption of a single CpG probe by a SNP linked to the lead SNP as the rate of change at multiple CpGs was associated with the lead SNP in many cases.

Overall, our analyses suggest that local SNPs can affect the rate at which DNA methylation changes with age at CpGs located in their vicinity.

### Local sequence context affects methylation trajectories with age

To understand the mechanistic basis of slope-QTLs, we analysed their genomic locations. CpGs that were part of slope-QTLs were significantly depleted from CpG islands and their shores but significantly enriched in intergenic regions and PMDs (35.4% and 27.6% of CpGs respectively, *Figure 4a*). Consistent with this, slope-QTL CpGs were also significantly enriched in chromatin states associated with enhancers and heterochromatin in GM12878 lymphoblastoid cells (12.3% and 39.1% of CpGs respectively, *Figure 4b*). Significant enrichments in enhancer and quiescent heterochromatin states were also seen in 95.7% and 100% of the 23 primary blood cell types analysed in the Roadmap Epigenomics project (p <0.05, *Supplementary Figure 4a*).

**Figure 4.**
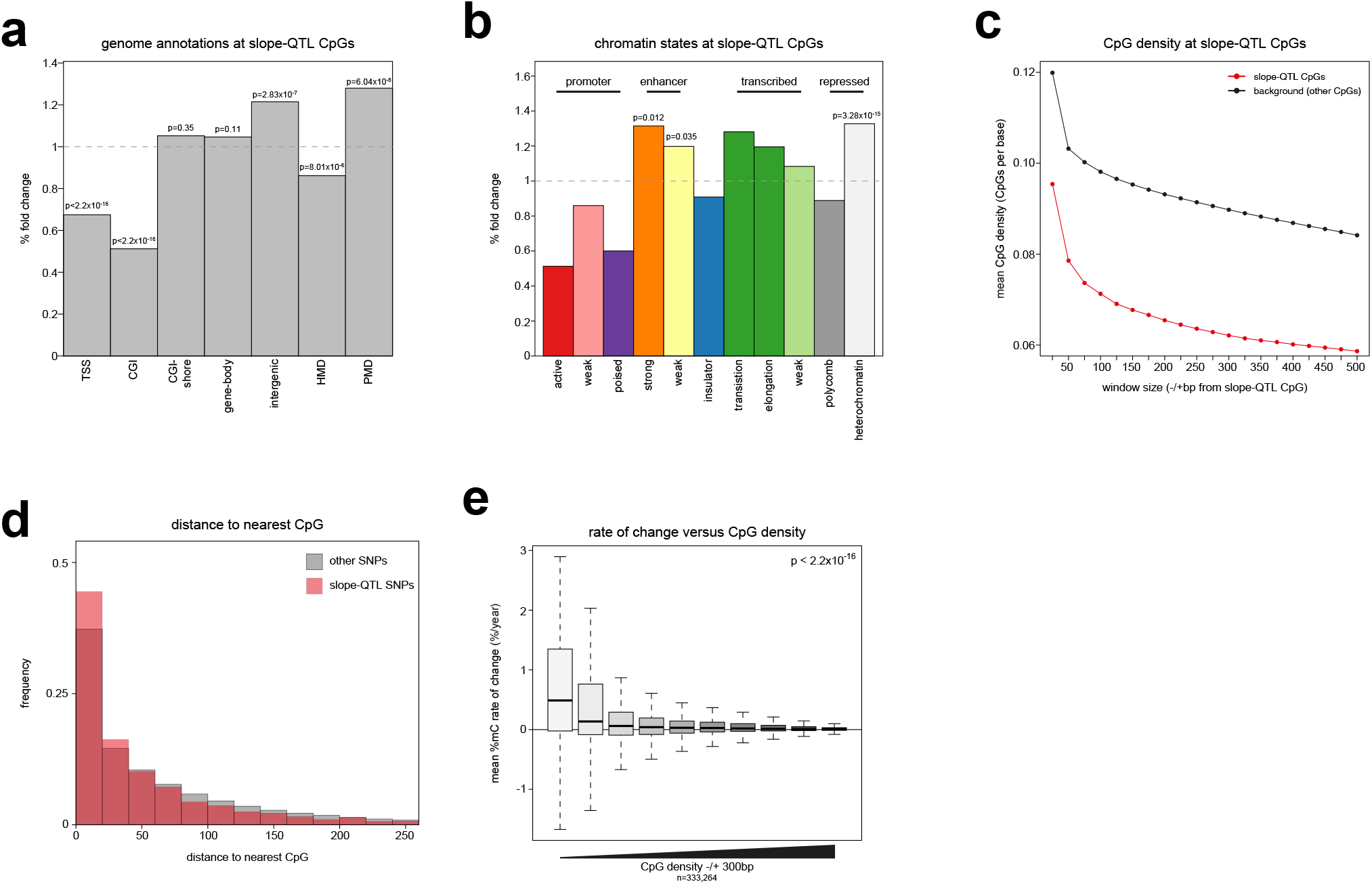
**a)** Slope-QTL CpGs are enriched in intergenic regions. Barplot showing the % fold change observed for slope-QTL CpGs in different genome annotations versus the background of all analysed CpGs. P-values are from 2-sided Fisher’s exact tests. **b)** Slope-QTL CpGs are enriched in enhancer and heterochromatin states in GM12878 cells. Barplot showing the % fold change observed for slope-QTL CpGs in different chromatin states in GM12878 cells versus the background of all analysed CpGs. Shown are significant P-values from 1-sided Fisher’s exact tests. **c)** Slope-QTL CpGs are found in regions of low CpG density. Line plot showing the mean CpG density around slope-QTL CpGs in different window sizes. Red shows slope-QTL CpGs and black shows all other CpGs assayed. **d)** Slope-QTL SNPs are found close to CpG sites. Histogram of the distances between slope-QTL SNPs and their nearest CpG site. Red shows the distribution for slope-QTL SNPs and grey shows all other SNPs assayed. **e)** CpG density is a major determinant of methylation trajectories with age. Boxplot showing mean methylation trajectories plotted against CpG density −/+ 300bp from the CpG. CpG density is binned into equally-sized groups. Lines=median; Box=25th–75th percentile; whiskers=1.5× interquartile range from box.

Previous work has highlighted local CpG density and the sequence surrounding intergenic CpGs as being associated with their methylation levels^15,39^. Given the enrichments we observed in intergenic annotations for slope-QTL CpGs and the observation that rapid gain CpGs have a low surrounding CpG density, we wondered if slope-QTLs might also be located in regions of low CpG density. Slope-QTL CpGs were located in regions with a significantly lower local genome CpG density than other CpGs assayed on the Infinium array (*Figure 4c*). The difference persisted across different window sizes surrounding the slope-QTL CpG although the effect was strongest around −/+ 325bp (*Figure 4c, Supplementary Figure 4b*). We then asked whether the SNPs associated with alterations in methylation trajectories at slope-QTLs might affect the local sequence composition around CpGs. To do so, we measured the distance between the slope-QTL lead SNP and its nearest CpG (irrespective of whether it is the Infinium array assayed slope-QTL CpG). Despite being located in CpG poor regions, slope-QTL lead SNPs were also found significantly closer to their closest CpG than non-slope-QTL SNPs in our analysis (T-test, p=5.4×10^−12^) and 11.4% directly affected a CpG site or the bases adjacent to one (159 out 1397).

These results suggest that alterations in the local sequence context around CpGs affects their methylation trajectory, particularly in regions of low CpG density. To understand the relationship between methylation trajectories and CpG density more generally, we analysed how methylation trajectories varied with local CpG density genome-wide. We observed that the magnitude of change at CpGs with age was significantly associated with their local CpG density (p < 2.2×10^−16^, T-test, *Figure 4e*). CpGs with a lower surrounding CpG density were more likely to have altered their methylation levels than those in higher CpG density regions and this was skewed towards gains of methylation. This effect was replicated in individuals aged over 65 in the STRADL cohort across the larger set of 742,122 CpGs assayed on the Illumina EPIC arrays (p < 2.2×10^−16^, T-test, *Supplementary Figure 4c*). The relationship between mean methylation trajectories and CpG density was also observed when we corrected for measured white blood cell counts from the LBC cohort (p < 2.2×10^−16^, T-test, *Supplementary Figure 4d*) demonstrating that it did not result from altered blood composition with age.

While the mean trajectories of CpGs located in low CpG density regions had a median gain of methylation with age, the mean trajectories were also far more variable in low CpG density regions (*Figure 4e*). We wondered whether this variability might also occur between individuals. To test this hypothesis, we calculated the variance in slope across individuals for each CpG. This slope variance displayed a strong parabolic relationship with the mean level of methylation at CpGs across the timepoints (*Supplementary Figure 4e*). After accounting for this relationship (see methods), we validated whether predicted differences in slope variance between CpGs could be observed. Given their utility in measuring age, CpGs which are part of the Hannum and Horvath epigenetic clocks would be expected to have consistent methylation trajectories and thus low inter-individual variance in slope. We found that this was the case and the slope variance of clock CpGs was significantly lower than other CpGs (*Supplementary Figure 4g*, Wilcoxon test, p=8.72×10^−6^). Analysing CpG slope variance more globally, we found a significant association between slope variance and local CpG density with CpGs in lower density regions having a greater slope variance (*Supplementary Figure 4f*, p < 2.2×10^−16^ by T-test).

Taken together, our genome-wide analysis therefore suggests that local CpG density is a strong determinant of a CpGs methylation trajectory with age and that changes in methylation with age are more likely to occur in regions of low CpG density. In addition, our results also suggest that alterations in a CpGs local sequence context caused by SNPs can alter its methylation trajectory with age.

## Discussion

While alterations in DNA methylation patterns with age have been widely observed in humans and are associated with health, the molecular mechanisms underpinning them remain unclear. Here we use human longitudinal DNA methylation profiles to demonstrate a strong effect of CpG density on methylation trajectories and that these can be altered by polymorphisms around CpGs (*Figure 5*).

**Figure 5.**
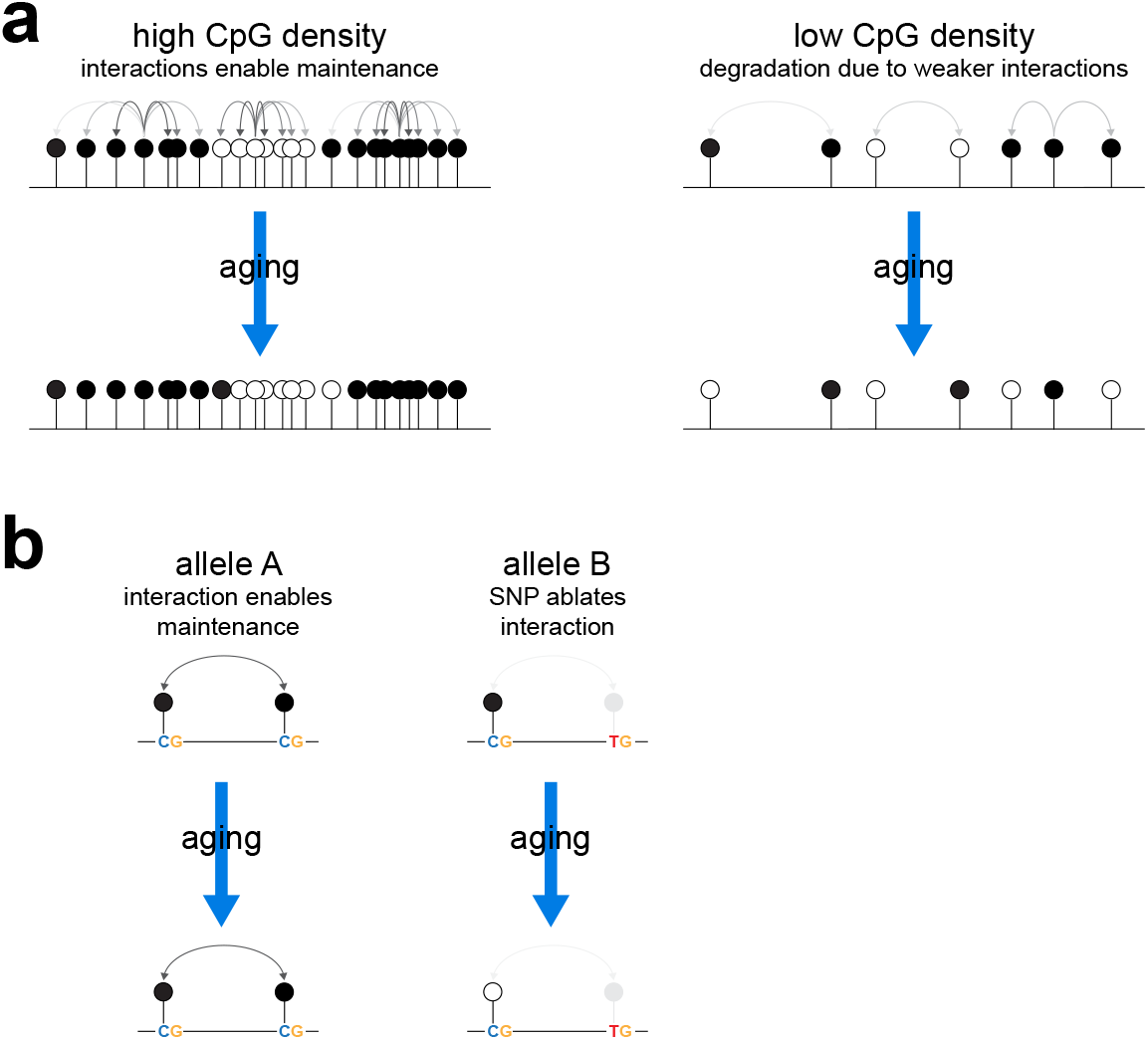
Local CpG density affects the trajectory of age-associated epigenetic changes. We propose that collaborative interactions between CpGs reinforces maintenance of methylation patterns in CpG dense regions (**a**). These interactions are weaker in CpG poor regions leading to the degradation of methylation patterns with time. These interactions can also be altered by SNPs leading to differences in epigenetic trajectories between individuals who inherit different alleles (**b**).

Previous work has described genetic influences on how DNA methylation patterns change with age. Genome-wide association studies (GWAS) find a number of loci that affect aging as estimated by epigenetic clocks^40–42^. In addition, analyses of rare Mendelian traits suggest that epigenetic aging is accelerated in two growth disorders, Sotos syndrome and Tatton-Rahman-Brown syndrome^43,44^. These two sets of studies analysed overall changes in the methylome with age rather than factors influencing the rate of change at individual loci. Further population analyses have also demonstrated the potential for genetic effects on how methylation changes with age at individual loci^23,24^ but did not define the mechanisms that underpin these associations. In contrast, an experiment analysing DNA methylation in mice possessing a copy of human chromosome 21 found that the introduced human loci changed their methylation status at a rate consistent with the mouse rather than human genome^25^. Based on these observations, the authors suggested that local sequence has little effect in determining the rate of change in DNA methylation with age. They suggested that this is instead primarily determined by the cellular environment. We demonstrate that local DNA sequence does play a role in age-associated methylation changes by showing that local SNPs can alter methylation trajectories and that local CpG density has a strong effect on the rate of change with age at individual CpGs in humans *in vivo*.

CpG density is known to be a determinant of steady state DNA methylation patterns in cells. The most highly CpG dense portions of the genome are CGIs which are typically DNA methylation free^45^. Variation in CpG density alone can also predict the methylation levels of bacterial DNA fragments integrated into the genome of mouse embryonic stem (mES) cells^46^. Analysis of Whole Genome Bisulfite Sequencing (WGBS) from human cell lines also revealed a strong relationship between CpG density and DNA methylation in heterochromatic PMDs independently of CGIs^39^. These are typically gene and CpG poor compared to euchromatic portions of the genome^34^. However, at individual CpGs within PMDs, methylation levels are positively correlated with the surrounding CpG density^39^. Analysis of diverse methylomes by WGBS shows that isolated CpGs have lower levels of methylation in PMDs in human tissues^15^. However, these observations are all derived from static snapshots of individual samples from cell lines and tissues. Here we provide the first demonstration of an effect of CpG density on the rate of change of methylation with age observed longitudinally *in vivo* in humans.

Interactions and collaborative reinforcement of DNA methylation between adjacent CpG sites has been proposed as being vital to maintain DNA methylation patterns by mathematical modelling^47^. Detailed analysis of methylation dynamics in mES cells possessing a sole DNMT, DNMT1, also finds evidence of neighbour-guided error correction as being important in maintaining DNA methylation patterns^48^. These collaborative interactions between CpGs are likely to be strongly influenced by the distance between CpGs. Thus, in lower CpG density regions, the greater distance between CpGs could result in weaker collaboration between neighbouring CpGs enabling greater degradation of developmental methylation patterns with age (*Figure 5*). These interactions are likely to be mediated by the molecular properties of the DNA methylation machinery. DNMT1 and DNMT3B both methylate processively along DNA strands whereas DNMT3A methylates in a distributive manner but can form multimers along the DNA fibre^49^. A computational analysis of DNA re-methylation dynamics suggests these processes are required to explain the observed rates of DNA re-methylation following replication^50^. The efficiency of DNMTs is also influenced by bases surrounding CpGs *in vitro^51,52^* and *in vivo^53^*. These preferences could affect the efficiency by which some CpGs are methylated and could explain the effect of some of the SNPs we have uncovered that do not directly affect CpGs. As well as being the target of DNMTs, unmethylated and methylated CpG nucleotides are also bound by (CXXC) and Methyl-Binding Domain (MBD) respectively^54–57^. It is possible that these proteins also mediate the effects of CpG density on DNA methylation changes with age. CXXC proteins include TET1 which plays a role in demethylation as well as CFP1, MLL1 and MLL2 which deposit the histone modification H3K4me3^58^. H3K4me3 inhibits the activity of de novo DNMTs through their ADD domains^59^. Thus, dense unmethylated CpGs can recruit proteins that reinforce their unmethylated status. It is unclear whether MBDs positively reinforce methylation patterns through binding to methylated CpGs, however, their binding in the genome tracks CpG density^60^.

Genetic effects on DNA methylation levels at individual CpGs independently of age have been widely documented in human populations as allele-specific methylation and meth-QTLs^19^. These are hypothesised to reflect the alteration of transcription factor binding by sequence polymorphisms with downstream effects on DNA methylation particularly at enhancers. TF binding can be both promoted or hindered by DNA methylation^61^. However, the majority of TF binding sites in the genome have reduced methylation^62^ and TFs have been strongly implicated in programming DNA methylation in cell lines^22^. The hypothesis that changes in TF binding underpin meth-QTLs is also supported by a study showing that many *trans* meth-QTLs correspond to TF genes^63^ and the analysis of a subset of SNPs affecting TF binding motifs in lymphoblastoid cell lines^64^. We find that the majority of slope-QTLs are in intergenic, heterochromatic regions with a low CpG density. These regions are depleted in both genes and enhancers^34^ suggesting slope-QTLs may be explained by alternative mechanisms. However, it is likely that slope-QTLs are not all explained by a single mechanism. 12.3% of slope-QTL CpGs are located in regions marked with enhancer-associated histone modifications in blood cells and these may be explained by TF dependent mechanisms.

In the current work, we have modelled methylation trajectories linearly. Previous studies of cultured fibroblasts^65^ and cross-sectional human cohorts^66^ suggest non-linear dynamics at some CpGs. Given the number of observations available per individual in the LBC cohort, non-linear trajectories cannot be fitted sufficiently robustly. We do observe that rapid gain CpGs gain methylation in the older individuals of LBC but show lower level of methylation with age in the younger members of the Generation Scotland cohort, an observation consistent with a non-linear trajectory over the lifecourse. Previous work has shown an overall loss of DNA methylation with age^5^. This is particularly prominent at low CpG density intergenic regions^15^. Our replicated observation of a set of heterochromatic, low CpG density loci that gain methylation later in life is therefore intriguing and warrants further investigation. However, we also find that low CpG density regions have a higher variability in their methylation trajectories suggesting that they might be better characterised by displaying greater epigenetic drift between individuals than a gain of methylation *per se*.

In addition to *cis*-slope-QTLs we also find evidence for the existence of *trans*-slope-QTLs. Previous work has documented large numbers of *trans*-meth-QTLs^20^ which have been ascribed to alterations in the expression of TFs or DNA methylation regulatory factors^63,67^. It is likely that more t*rans*-slope QTLs exist. However, due to the large multiple testing burden incurred by searching every CpG against every possible SNP, it was not possible to detect large numbers of them in the current study. Future investigation of *trans*-slope-QTLs will require larger studies with a higher statistical power in combination with approaches which reduce the multiple testing burden by taking account of the lack of independence between SNPs^68^.

Taken together our study suggests that DNA sequence, and CpG density in particular, is a major influence on the local tick rate of age-associated DNA methylation changes. We ascribe this effect to interactions between neighbouring CpGs reinforcing maintenance of methylation patterns through the action of the DNA methylation machinery.

## Materials and Methods

### Statistical analysis

Statistical testing was performed using *R v4.0.2*. All tests were two-sided, unless otherwise stated. All linear models were fitted using the *lme4* package (*v1.1-21*). Further details of specific analyses provided in the relevant methods sections below.

### Cohort details

The Lothian Birth Cohort 1936^26–28^ is derived from a set of individuals born in 1936 who had mostly taken part in the Scottish Mental Survey 1947 at a mean age of 11 years as part of national testing of almost all children born in 1936 who attended Scottish schools on 4 June 1947. A total of 1,091 participants who were living in the Lothian area of Scotland were re-contacted in later life. DNA methylation was measured for this cohort around 70 years of age and subsequently at a mean of 73, 76 and 79 years on Illumina 450k arrays. In total this corresponds to 2852 samples from 1056 unique individuals. In this study we focused on the 600 individuals for whom ≥ 3 methylation measurements existed (283 female and 317 male). A breakdown of the sample demographics can be found in *Table 1*.

The Generation Scotland dataset was derived from a subset of individuals in the Generation Scotland or Scottish Family Health Study (GS:SFHS) cohort. GS:SFHS is a family-based population cohort investigating the genetics of health and disease in approximately 24,000 individuals across Scotland^32^. Baseline data were collected between 2006 and 2011. The subset used here focuses on 5,101 individuals aged 18-95 years for whom Illumina EPIC array data had been collected from blood at baseline contact^31^.

All participants provided written informed consent. Ethical permission for the Lothian Birth Cohort 1936 study protocol was obtained from the Multi-Centre Research Ethics Committee for Scotland (Wave 1: MREC/01/0/56), the Lothian Research Ethics Committee (Wave 1: LREC/2003/2/29), and the Scotland A Research Ethics Committee (Wave 2-4: 07/MRE00/58). Ethical consent for GS:SFHS was similarly granted at the initiation of the study (05/S1401/89) and for subsequent follow-on study (14/SS/0039).

### Processing of Illumina Infinium array data

Infinium arrays from both the LBC and STRADL cohorts were processed from IDAT files. These were normalised using the *ssNoob* method from the Bioconductor package *minfi* (*v1.22.1*) to derive beta values and detection p-values (beta threshold = 0.001)^69,70^. Individual beta values were excluded where detection p-value was > 0.01. Non-CG probes and probes not located on autosomes were also excluded from the analysis. Infinium probe locations in the hg38 genome build were taken from Zhou *et al*^29^. Probes categorised as overlapping common SNPs or having ambiguous genome mapping in that paper were excluded from the analysis (*MASK.snp5.common, MASK.mapping, MASK.sub30.copy* from Zhou *et al*)^29^. Throughout the paper, beta values are converted to estimated % methylation measurements by multiplying by 100.

### Processing of SNP data

DNA samples from the Lothian Birth Cohort 1936 were genotyped at the Wellcome Trust Clinical Research Facility using the Illumina 610-Quadv1 array (San Diego)^71^. Individuals were excluded based on relatedness (n = 8), unresolved sex discrepancy (n = 12), low call rate (≤0.95 n = 16) and evidence of non-European descent (n = 1). SNPs were included if they had a call rate ≥0.98, a minor allele frequency ≥0.01, and a Hardy-Weinberg equilibrium test with p ≥ 0.001.

### Modelling of DNA methylation trajectories

DNA methylation trajectories for each CpG in each individual were modelled by fitting a linear model of beta value on age using *R* and the *lme4* package. This was only done for individuals and CpGs for which ≥ 3 datapoints were present in the processed dataset. Slopes for each individual were taken from the linear models. CpGs were considered to have a rate of change significantly different from 0 if the distribution of slopes for all individuals had a Bonferroni corrected p-value < 0.01 by T-test.

We defined rapid gain CpGs as those with significant slopes where the mean change in Beta per year was greater than the local minimum on a histogram of all CpGs with a significant slope (bin size = 0.0005 and threshold > 0.0159, both in beta/year).

When modelling methylation trajectories in the Generation Scotland dataset, linear models of beta on age were calculated across all individuals present in the analysis and slope coefficients extracted. To compare individuals of a similar age to those in LBC we used only the 406 individuals aged >= 65 years from Generation Scotland where indicated.

To correct for variation in blood cell populations we used white blood cell counts (neutrophils, lymphocytes, monocytes, eosinophils and basophils) for each sample collected on a Sysmex HST system under standard operating procedures within the National Health Service Haemotology laboratory, Western General Hospital, Edinburgh. We then used these to derive residualised beta values corrected for variation in these blood cell populations by fitting a linear model:

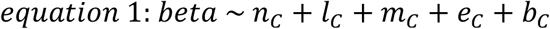

Where: *n_c_* = neutrophil count, *l_c_* = lymphocyte count, *m_c_* = monocyte count, *e_c_* = eosinophil count and *b_c_* = basophil count. The residuals of this model where then used as corrected beta values and methylation trajectories were then modelled from them as above.

The variability in methylation trajectories at each CpG was investigated by calculating the variance of all individual linear model slope coefficients for that CpG. As the variance was strongly related to the mean beta value of each CpG (*Supplementary Figure 4F*), we normalised variance of CpGs by calculating the median of 20 equal bins based upon the mean beta and then subtracting the calculated median from all the CpGs in that bin. These residualised variances were then analysed.

### Analysis of epigenetic clocks

Horvath epigenetic clock CpGs were taken from Horvath 2013 *Supplementary Table 3*^6^. This table contains details of the age relationship these CpGs displayed in the original derivation of this clock^6^. We defined Horvath clock increasing and decreasing CpGs from these coefficients. Hannnum epigenetic clock CpGs were taken from Hannum *et al* 2013 *Supplementary Table 3*^7^. As this did not include details of the rate of change at each clock CpG over time, we determined increasing and decreasing CpGs using the STRADL cross-sectional dataset by fitting linear models to each as described above.

### Analysis of CpG annotation

CpGs were annotated to CGIs based on Illingworth *et al*^72^. Overlapping CGI intervals were merged using BEDtools (*v2.27.1*)^73^ before they were converted to hg38 positions using the UCSC browser *liftover* tool. CpGs were then overlapped with CGIs using BEDtools. CGI shores were defined as the 2Kb on either side of each CGI using BEDtools and similarly overlapped with CpGs. CpGs were annotated relative to genes using BEDTools to overlap them with ENSEMBL protein coding genes (Ensembl Release 98/GCRh38). CpGs were annotated as being located at a transcription start site (TSS) if they overlapped a protein coding TSS and as located in a gene body if they overlapped a transcript but not a TSS. The remaining CpGs which did not overlap a TSS or transcript were annotated as intergenic. PMD and highly methylated region (HMD) definitions were taken from Zhou *et al*^15^ from *commonPMDs* and *commonHMDs* defined across 40 tumour and 9 normal samples and overlapped with CpGs using BEDtools.

Infinium probes were mapped to existing ChromHMM annotations^74^ using the BEDtools *intersect* function (*v2.27.1*)^73^. Identical ChromHMM labels were merged for analysis. To test for enrichment of an annotation, a Fisher’s exact test was performed for number of rapid gain CpGs or slope-QTL CpGs against number of control probes. ENCODE GM12878 ChromHMM^36^ annotations were downloaded as bedfiles from the UCSC genome browser. Roadmap Epigenomics ChromHMM annotations for primary human cell types were downloaded as mnemonics BED files from the Roadmap Epigenomics site (*https://egg2.wustl.edu/roadmap/data/byFileType/chromhmmSegmentations/ChmmModels/coreMarks/jointModel/final*)^37^. The 23 primary blood cell types analysed were selected by manual examination of Roadmap Epigenomics sample annotations.

To calculate local CpG density, windows of sequence (eg −/+ 300bp) were extracted around each CpG analysed from the hg38 genome sequence using BEDtools (*v2.23.0*) and the number of CpG dyads within this window counted.

### Slope-QTL analysis

Associations between genotype and local rates of methylation change (quantified as the slope coefficient for each individual linear model) were analysed using the *matrix-eQTL R* package (*v2.23*)^75^. For *cis*-associations, we set a distance cut-off of 1Mb. Significant associations were those where the Benjamini-Hochberg corrected p-value as < 0.05. The analysis of *trans*-associations was carried out similarly with the distance threshold removed. To remove loci where apparent differences in the rate of change associated with genotype were caused by effects on the variability of the slope, we tested for associations between SNPs within 1Mb and the variance of methylation at each CpG (quantified as the residual sum of each individual linear model). CpG SNP pairs with a significant association to variability were defined as those with a matrix-eQTL Benjamini-Hochberg corrected p < 0.05.

We then performed a conditional analysis to determine how many independent SNP-CpG pairs were present. The SNP with the most significant p-value associated with each CpG was designated the lead SNP, and then all other associated SNPs were tested against the residuals of the linear model of the lead SNP. Other SNPs were then designated as independent hits if their p-value of association was < 0.05 following Bonferroni correction.

### Age x genotype interaction modelling

A linear model (methylation~ age*genotype) was fitted to each CpG in LBC. The correlation between these Age x Genotype effect sizes and the rate of methylation change~genotype effect size in the LBC was calculated using Pearson’s coefficient.

## Data availability

According to the terms of consent for Lothian Birth Cohort 1936 data are available on request from the Lothian Birth Cohort Study, University of Edinburgh. Similarly, according to the terms of consent for Generation Scotland participants, access to data must be reviewed by the Generation Scotland Access Committee. Applications should be made to.

## Acknowledgements

We thank Riccardo Marioni, Chris Haley, Ailith Ewing, David Porteous, Chris Ponting, Rob Illingworth, Tamir Chandra, Sara Hagg, Yunzhang Wang and Chantriolnt-Andreas Kapourani and Nick Gilbert for helpful discussions about the study and the manuscript. This work has made use of the resources provided by the University of Edinburgh digital research services and the MRC IGMM compute cluster. DS is a Cancer Research UK Career Development fellow (reference C47648/A20837), and work in his laboratory is also supported by an MRC university grant to the MRC Human Genetics Unit. S.R.C. and I.J.D. were supported by a National Institutes of Health (NIH) research grant R01AG054628, and S.R.C is supported by a Sir Henry Dale Fellowship jointly funded by the Wellcome Trust and the Royal Society (221890/Z/20/Z). AMM is supported by the Wellcome Trust (104036/Z/14/Z, 216767/Z/19/Z, 220857/Z/20/Z) and UKRI MRC (MC_PC_17209, MR/S035818/1). PMV acknowledges support from the Australian National Health and Medical Research Council (1113400) and the Australian Research Council (FL180100072). DMH is supported by a Sir Henry Wellcome Postdoctoral Fellowship (Reference 213674/Z/18/Z). We thank the LBC1936 participants and team members who contributed to the study. Further study information can be found at https://www.ed.ac.uk/lothian-birth-cohorts. The LBC1936 is supported by Age UK (Disconnected Mind project, which supports S.E.H.), the Medical Research Council (G0701120, G1001245, MR/M013111/1, MR/R024065/1), and the University of Edinburgh. Genotyping of LBC1936 was funded by the BBSRC (BB/F019394/1), and methylation typing of LBC1936 was supported by Centre for Cognitive Ageing and Cognitive Epidemiology (Pilot Fund award), Age UK, The Wellcome Trust Institutional Strategic Support Fund, The University of Edinburgh, and The University of Queensland. We are grateful to all the families who took part in the Generation Scotland study along with the general practitioners and the Scottish School of Primary Care for their help in recruiting them, and the entire Generation Scotland team, which includes interviewers, computer and laboratory technicians, clerical workers, research scientists, volunteers, managers, receptionists, healthcare assistants, and nurses. Work on Generation Scotland was supported by a Wellcome Strategic Award “STratifying Resilience and Depression Longitudinally” (STRADL; 104036/Z/14/Z) to AMM, KLE, and others, and an MRC Mental Health Data Pathfinder Grant (MC_PC_17209) to AMM. Generation Scotland received core support from the Chief Scientist Office of the Scottish Government Health Directorates (CZD/16/6) and the Scottish Funding Council (HR03006). DNA methylation profiling and analysis of the GS:SFHS samples was supported by Wellcome Investigator Award 220857/Z/20/Z and Grant 104036/Z/14/Z (PI: AM McIntosh) and through funding from NARSAD (Ref: 27404; awardee: Dr DM Howard) and the Royal College of Physicians of Edinburgh (Sim Fellowship; Awardee: Dr HC Whalley).

## Contributions

JH and DS conducted the computational analysis presented in the manuscript. RMW, SEH, SRC, DMH, and ELH curated data relating to the study. QC and PMV contributed to the analysis and interpretation of results. ALS, JDS, GDW, ADM, KLM, AMM, IJD ascertained subjects, obtained samples and funding for the profiling of cohort samples. DS planned and supervised the study and wrote the manuscript with input from JH and review by all authors.

