## Supplemental figures and legends for "Local CpG density affects the trajectory of age-associated epigenetic changes"

**Supplementary Figure Legends**

**Supplementary Figure 1**

**a)** Mean methylation trajectories calculated from individual longitudinal data are highly correlated with the rate of change calculated from cross-sectional data. Density scatter plot of the mean methylation trajectory against the rate of change estimated from the entire population.

**b)** Slopes at epigenetic clock CpGs calculated from individual methylation trajectories are reproduced in a cross-sectional dataset. Scatter plot comparing the slopes calculated from the longitudinal LBC dataset to those calculated in the cross-sectional Generation Scotland dataset. Line indicates identity.

**c)** Epigenetic clock CpGs show modest slopes. Histogram of the mean slope calculated from individual methylation trajectories for CpGs that are part of epigenetic clocks (red) and all other assayed CpGs (grey).

**Supplementary Figure 2**

**a)** Rapid gain CpGs show lower methylation with age in earlier life. Boxplot of the mean methylation levels observed at rapid gain CpGs for individuals in the STRADL cross-sectional cohort grouped by age. Lines=median; Box=25th–75th percentile; whiskers=1.5× interquartile range from box.

**b)** Rapid gain CpGs gain methylation after correction for variation in blood cell counts. Histogram of the rate of change in DNA methylation (corrected beta values/year) calculated from the LBC cohort after correction for measured white blood cell type counts. Shown are rapid gain CpGs (red) and all other CpGs (grey).

**c)** Rapid gain CpGs are found in regions of low CpG density. Boxplot comparing the CpG density -/+ 300bp from the CpG at rapid gain CpGs to all CpGs assayed. Lines=median; Box=25th–75th percentile; whiskers=1.5× interquartile range from box. P-value was calculated by Wilcoxon rank sum test.

**d)** Rapid gain CpGs are enriched in transcription and heterochromatin states in primary blood cells. Heatmap of the odds-ratios of rapid gain CpGs being enriched in different chromatin states in primary blood cells compared to the background of all assayed CpGs. Odds ratios derived from Fisher's exact tests.

**Supplementary Figure 3**

**a)** Effect sizes of slope-QTLs are significantly correlated with those calculated from age x genotype models. Scatter plots of the effect sizes calculated for slope-QTLs against the age x genotype effect calculated for the same CpG-SNP pair.

**Supplementary Figure 4**

**a)** Slope-QTL CpGs are enriched in enhancer and heterochromatin states in primary blood cells. Heatmap of the odds-ratios of slope-QTL CpGs being enriched in different chromatin states in primary blood cells compared to the background of all assayed CpGs. Odds ratios derived from Fisher's exact tests.

**b)** CpG density differences around slope-QTLs are greatest around 300bp from the affected CpG. Line plots showing the ratio (left) and p-value(right) of differences in the CpG density at different window sizes around slope-QTL CpGs and the background of all assayed CpGs. P-values were calculated using a T-test.

**c)** CpG density is a major determinant of methylation trajectories with age in a second independent cohort. Boxplot showing estimated rates of change in DNA methylation from the generation Scotland cohort plotted against CpG density -/+ 300bp from the CpG. CpG density is binned into equally sized groups. Lines=median; Box=25th–75th percentile; whiskers=1.5× interquartile range from box.

**d)** The effect of CpG density on methylation trajectories is observed after correction for varying blood cell counts. Boxplot showing estimated rates of change in DNA methylation from the LBC cohort after correction for measured white blood cell counts plotted against CpG density -/+ 300bp from the CpG. CpG density is binned into equally sized groups. Lines=median; Box=25th–75th percentile; whiskers=1.5× interquartile range from box.

**e)** Variance in slope is strongly related to mean methylation level. Density scatter plot of the variance in slope observed for CpGs versus their mean methylation level across all timepoints in LBC.

**f)** Epigenetic clock CpGs show low variance in slope. Boxplot comparing the residualised variance in slope for CpGs that are part of the Hannum and Horvath epigenetic clocks and all other CpGs. Lines=median; Box=25th–75th percentile; whiskers=1.5× interquartile range from box; n= 337 and 294,518 for clock and other CpGs respectively.

**g)** Low CpG density regions display more variable methylation trajectories. Boxplot showing the variation in CpG slope across individuals from the LBC cohort plotted against CpG density -/+ 300bp from the CpG. CpG density is binned into equally sized groups. Lines=median; Box=25th–75th percentile; whiskers=1.5× interquartile range from box.

### Supplementary Figure 1

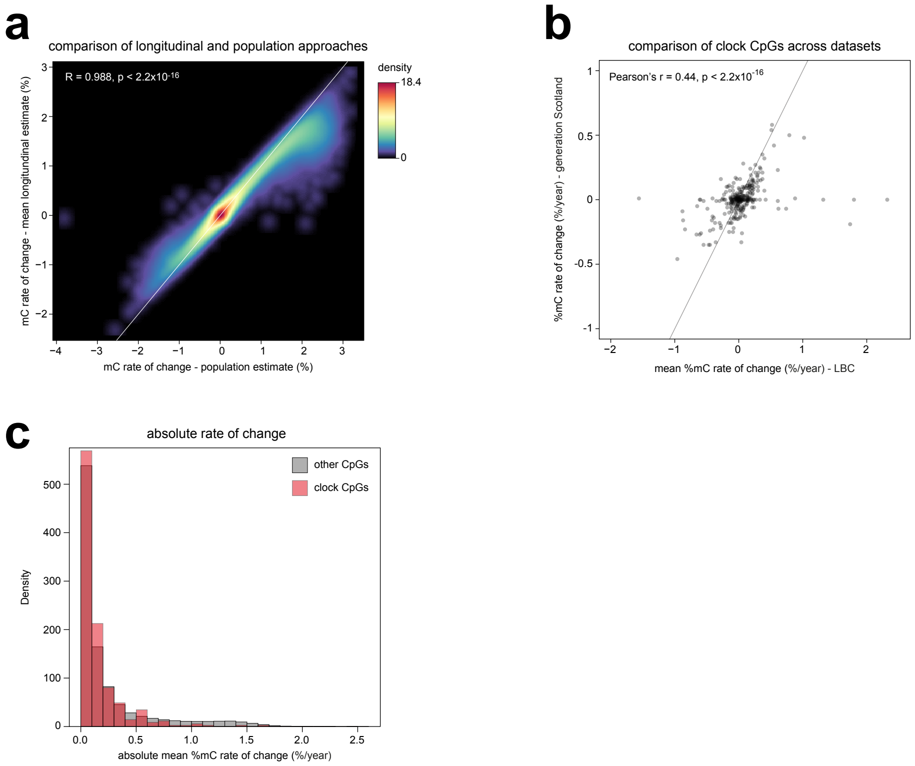

### Supplementary Figure 2

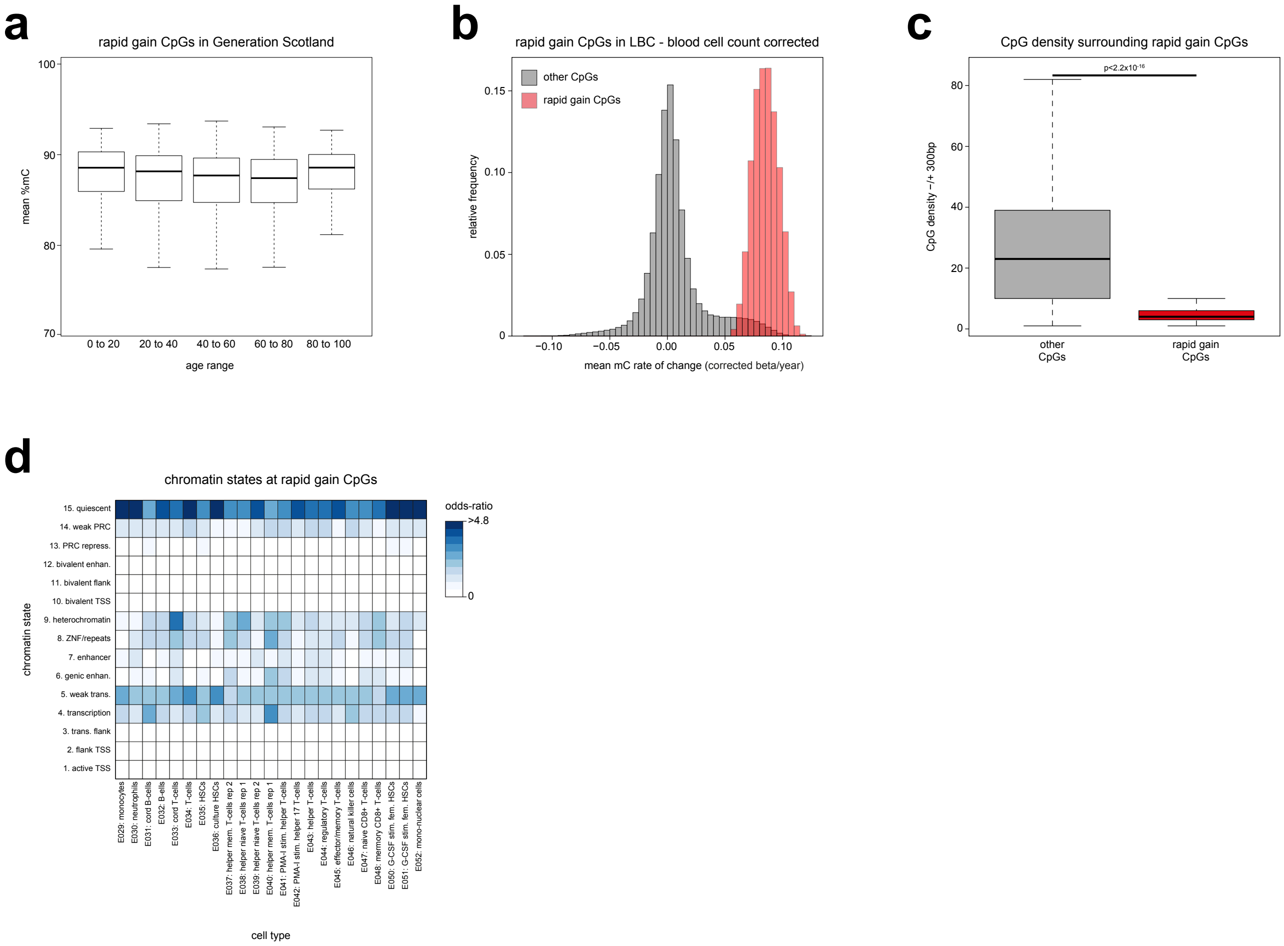

### Supplementary Figure 3

**a**

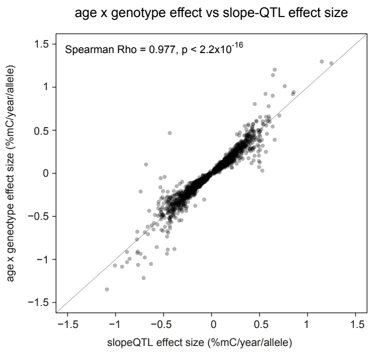

### Supplementary Figure 4

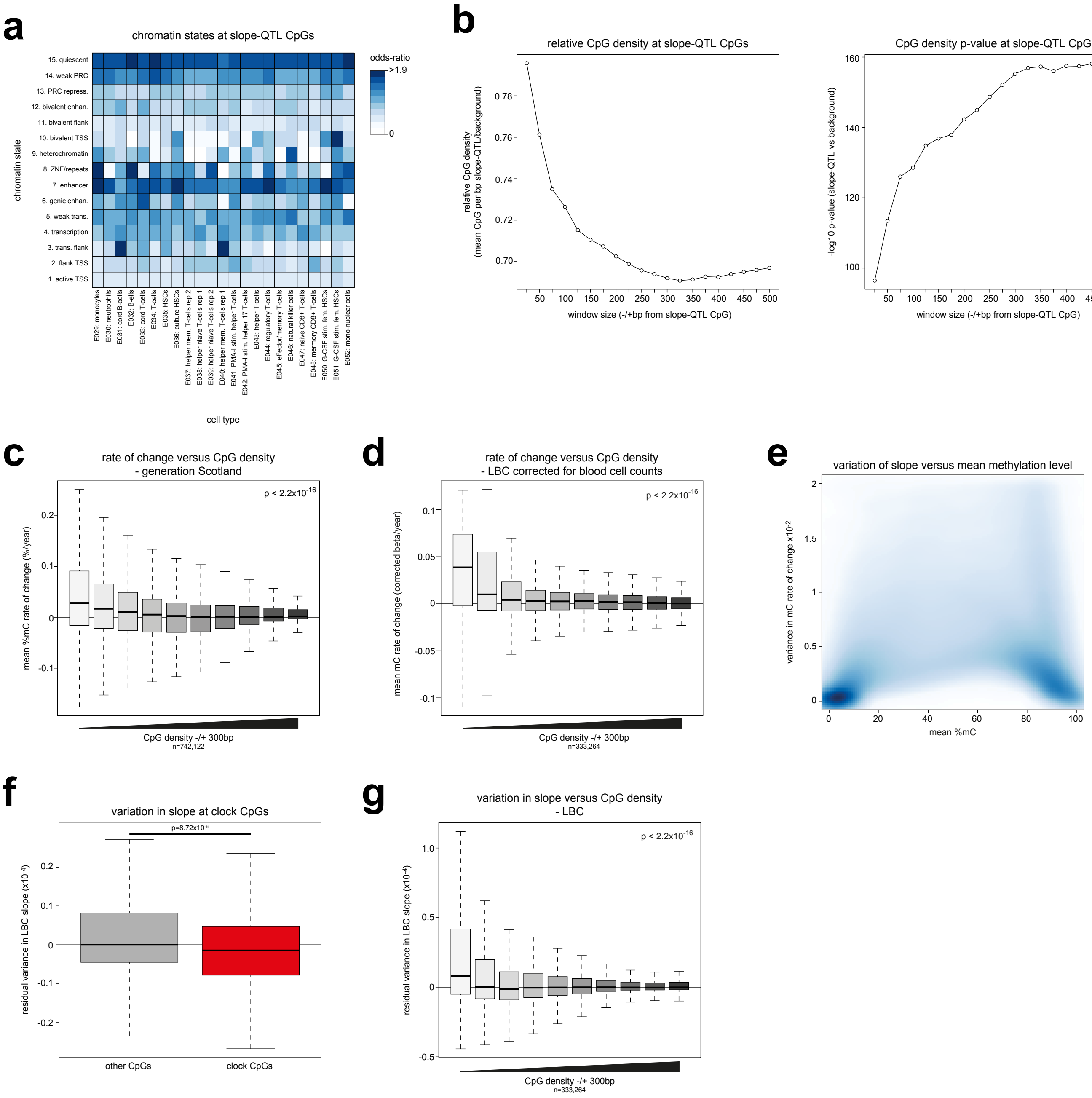
